# An effective workshop on “How to be an Effective Mentor for Underrepresented STEM Trainees”

**DOI:** 10.1101/2021.12.06.471498

**Authors:** Andrea G. Marshall, Caroline B. Palavicino-Maggio, Elsie Spencer, Zer Vue, Heather Beasley, Edgar Garza-Lopez, Lillian Brady, Zachary Conley, Kit Neikirk, Sandra Murray, Denise Martinez, Haysetta Shuler, Derrick Morton, Antentor Hinton

## Abstract

Despite an increase in programming to promote persons excluded by their ethnicity or race (PEER) scholars, minorities remain underrepresented in many STEM programs. The academic pipeline is largely leaky for underrepresented minority (URM) scholars due to a lack of effective mentorship. Many URM students experience microaggressions and discrimination from their mentors due to a lack of quality mentorship training. In this workshop, we provide a framework for how to be an effective mentor to URM trainees. Mentees, especially URM trainees, can flourish in effective mentoring environments where they feel welcomed and can comfortably develop new ideas without feeling threatened by external factors. Effective mentoring environments provide motivational support, empathy, cultural competency, and training.

## The framework of the workshop

Designing an effective mentorship workshop requires examples of the characteristics of effective mentors of underrepresented minority (URM) trainees, a blueprint for selecting mentees based on the mentorship environment, and strategies for maintaining nurturing mentor-mentee relationships. Mentor-mentee relationships require navigating unique challenges to ensure the success of URM trainees. Frequently, URM students face microaggressions, imposter fear, and difficulties in building networks and mentorship relationships, which increase the risk of falling out of the academic pipeline (Hinton Jr *et al*. 2020a; National Academies of Sciences 2020; Shuler *et al*. 2021; Uddin and De Los Reyes 2021). Despite the nature of these challenges, based on scientific findings, URM trainees perform better under supportive mentorship relationships, which can help them overcome the daily challenges that deters them from staying in the academic pipeline (Hinton Jr *et al*. 2020a; Termini *et al*. 2021a).

During the workshop strategies were presented to help potential mentors identify best practices for effective and motivational mentoring. The importance of celebrating the mentee and their journey was emphasized in the workshop, in addition the mentor was provided strategies for identifying mentees in need of special guidance. These strategies encouraged mentors to reframe how they think about students, work to have a more authentic mentor-mentee relationships, and develop a motivation-based mentoring approach based on emotional intelligence (Gardenswartz, Cherbosque and Rowe 2010; Opengart and Bierema 2015; Montgomery 2017; Hinton *et al*. 2020; Hinton Jr *et al*. 2020b; Shuler *et al*. 2021).

In the workshop, all participants were presented with a model of intentional mentorship. This mentoring model emphasizes a willingness to learn and establish credibility while facilitating the formation of positive relationships through networking (Shuler *et al*. 2021). Workshop participants were also exposed to effective and ineffective mentorship practices (Neikirk 2021). Furthermore, the workshop discussed using motivational mentoring as a way to cultivate the mentee’s spirit of excellence as they navigate their career development -- through the use of individual development plans (IDP) (National Academies of Sciences 2020; Shuler *et al*. 2021). These strategies are especially useful for mentees during times of hardship, such as classroom challenges. (McReynolds *et al*. 2020; Termini *et al*. 2021b).

## Introduction

Mentoring relationships are essential for the development of a mentee’s career, especially those from URM groups. Successful mentoring requires environments that bolster motivational ambition, provides empathy, and utilizes cultural competency. Toxic mentoring environments arise from poor communication, lack of commitment and experience, conflicting personalities, perceived competition, poor perceived performance, and difficulty in forming interpersonal connections with the mentee.

### Key goals for successfully mentoring diverse trainees

- Practice meditation and mindfulness
- Be aware of your own biases
- Create a positive environment
- Respond, do not react
- Promote self-motivation, encompass your inner resources to act, reach, and achieve goals and aspirations
- Provide constructive, not deconstructive, criticisms
- Cultural competence

### Motivational Support: How to motivate and support diverse trainees simultaneouslyã

Motivation is the inner drive to excel, which is often changed by internal and/or external conflict. This inner drive is important for cultivating goals and providing direction. A trainee’s motivation governs the direction of the trainee’s behavior in any mentoring environment, such as their effort, grit, and attitude. Minority trainees not only face external barriers during their educational and career journey, such as toxic mentorship and institutional inequities, but also experience internal challenges, such as John Henryism (Rolle *et al*. 2021). Furthermore, URM also face barriers including imposter fear, also known as imposter syndrome which is discussed in the workshop as a stigmatizing word that places the issue on the individual as opposed to the environment (Hinton Jr *et al*. 2020b; Rolle *et al*. 2021). These barriers stimulate a lack of confidence, which affects their perseverance. Since all trainees have different motivations, mentors should personalize their mentorship approach based on their mentee’s goals and motivations. A single mentorship strategy is often insufficient for a diverse group of mentees.

Mentees differ physically and emotionally, including in their motor, moral, and learning abilities. Mentors set an example for their trainees. Quality mentoring comes from being an inspiration (Shuler *et al*. 2021). Mentorship is an investment, not only to their institutions but to society as their mentees may make substantial contributions (Hinton Jr *et al*. 2020b; Shuler *et al*. 2021). Thus, the mentor’s character can play a big role in how mentees view themselves. An inspiring mentor listens, serves, shares, focuses on positivity, stays authentic, is willing to learn, and remains humble. Minority trainees excel with positive reinforcement. Mentors foster positivity in their relationships with mentees. Mentors must also identify and restrain negative beliefs that may influence their guidance and be willing to accept constructive feedback.

### Providing support and empathy

Mentors need to invest time in getting to know their mentees. Active listening is as important as intentional mentoring. An effective mentor sets aside time to speak with their mentees and pays attention to what they have to say (Shuler *et al*. 2021). Active listening, instead of passive listening, entails action. For example, if your mentee makes you aware of a concern or question, the mentor might not have an answer to a specific situation, which would require seeking out advice from their network. Intentional listening is essential to effectively communicate with mentees. For example, if a mentor does not completely understand their trainees’ questions or concerns, the mentor may consider asking for clarification, which ensures a clear line of communication between the mentor and mentee (Shuler *et al*. 2021). Active listening also requires avoiding distractions, such as emails, while providing their mentees with undivided attention. Focus on clarifying the situation to best provide an answer or suggestion. It is also important a mentor maintains an open mind. Obstacles and setbacks are a good way for mentees and mentors to grow and develop their skills.

Furthermore, each challenge is unique. Although mentees may be of the same gender, racial/ethnic background, socio-economic background, or school systems, they are all individuals with different journeys and motivations. No racial/ethnic group is monochromic hence it is essential to develop a personalized ‘individual development plan’ (IDP) (Hinton Jr *et al*. 2020b). Effective mentors also seek to maintain transparency by breaking down communication barriers, including seeking out alternative approaches, media, or technology to carry out conversations.

It is also important for mentors to focus on developing their emotional intelligence (EI), which is also known as emotional quotient or emotional intelligence quotient (Hinton Jr *et al*. 2020b, Shuler *et al*. 2021). EI is the ability of understanding feelings, emotional language, and signals conveyed by emotions (Hinton Jr *et al*. 2020b, Shuler *et al*. 2021).. EI involves distinguishing and managing our personal feelings and interactions from those of other people (Hinton Jr *et al*. 2020b, Shuler *et al*. 2021). EI assists with managing your behavior, navigating social areas, and helping others make critical life choices (Hinton Jr *et al*. 2020b, Shuler *et al*. 2021). EI also helps identify personal biases in thinking (Hinton Jr *et al*. 2020b, Shuler *et al*. 2021). It is like a window to determine why a mentee or colleague behaves a certain way or avoids making certain decisions. Practicing empathy towards their mentees can help mentors and mentees communicate more efficiently.

### Cultural competency and training

Cultural competence is the knowledge and skills needed to work with a diverse group in a meaningful relevant and productive way. Cultural competence involves an understanding of the role of religion, community, and culture in the lives and careers of underrepresented minority mentees. Mentors should familiarize themselves with common racial insensitivities and develop methods to ask questions on these topics with sensitivity & avoid perpetuating racial macro- and microaggressions.

Based on these concepts, we tested how students perceived the information and whether they could apply it to their career development and individual development plan. In this particular questionnaire, we used four questions to gauge interest. The questions consisted of a 10-point scale that was based on rating the following concepts: overall presentation, support team, verbal and nonverbal communication skills, and networking.

## Methods

Twenty-four students from Winston-Salem State University (a historically Black public university) attended a 90-minute virtual workshop. The participants completed an anonymous questionnaire before and after the workshop to gauge their expectations and satisfaction regarding the workshop (Table 1). The data were compared using nonparametric Wilcoxon matched-pairs and signed-rank tests to determine differences between measures. Differences were considered statistically significant when P values were less than 0.05. ****P < 0.0001; ***P < 0.001; **P < 0.01; *P < 0.05; NS, not significant P > 0.05; NS (not significant).

**Table 1.** Pre- and post-workshop evaluations.

| **Pre-workshop survey questions** | **Post-workshop survey questions** |
| --- | --- |
| On a scale of 1 to 10, do you think the presentation will keep you well informed? | On a scale of 1 to 10, how did you like the presentation? |
| On a scale of 1 to 10, how do you think the talk will improve your verbal and non-verbal communication? | On a scale of 1 to 10, how do you think the talk helped you improve your verbal and non-verbal communication? |
| On a scale of 1 to 10, how well do you think the talk will improve your networking skills? | On a scale of 1 to 10, how much do you think the talk helped you improve your networking skills? |
| On a scale of 1 to 10, how do you think the talk will improve your understanding of what a support team does? | On a scale of 1 to 10, how much do you think the talk helped you improve your understanding of what a support team does? |

## Results

We summarized the data from the questionnaires using box and whisker plots in which the red centerline denotes the median, and error bars denote the standard error. Individual values are represented by circles. Overall, participant feedback was positive. Responses to the pre-workshop questionaries suggest that mentees did not initially believe the workshop would be beneficial (Figure 1A-D, Pre-Test). The data suggests that their low expectations may be a result of low exposure or lack of mentorship.

**Figure 1.**
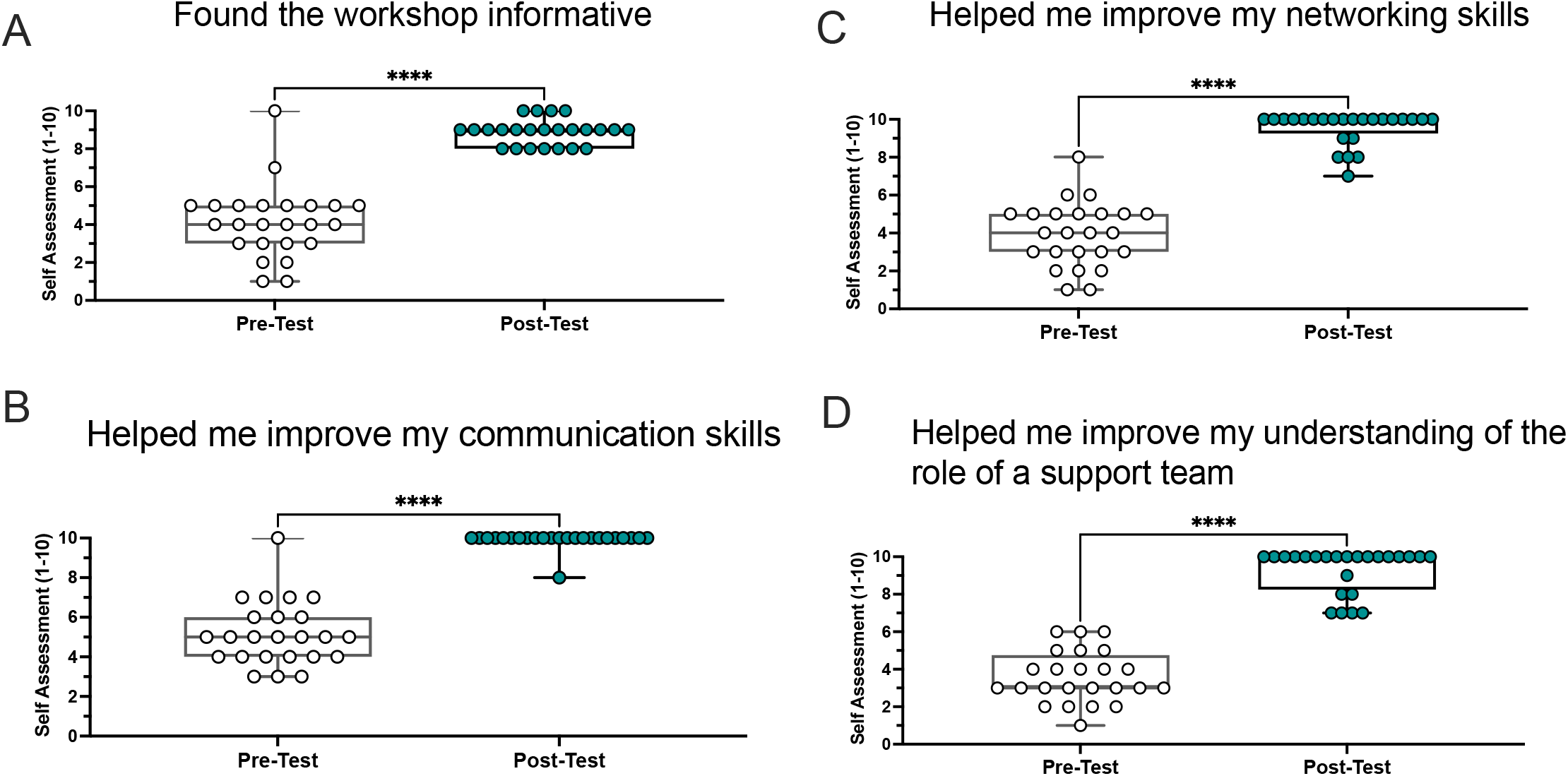
Results from pre- and post-workshop evaluations. These pre- and post-workshop questions were also used to evaluate mentees’ knowledge regarding mentee-mentor relationships. **A**. The informativeness of the workshop. **B**. How much the workshop improved communication skills. **C**. How much the workshop improved networking skills. **D**. How much the workshop improved understanding of support teams and assistive roles.

Importantly, inconsistent mentorship may alter the mentee trajectories (Packard 2003; Thomas, Willis and Davis 2007; Janis and Barker 2016). However, after the workshop, feedback scores increased by an average of 5.2 points on a 10-point scale. The median score was a 9 or higher for every question asked (Figure 1A-D, Post-Test), indicating that the workshop was found favorable and helpful for identifying mentors or considering mentors for other parts of their lives.

All post-workshop questionaries show a significant difference compared with pre-workshop questions. Initially, on average, participants believed the workshop to be low to moderately informative with an average evaluation of 4.1 (Figure 1A, Pre-Test). Following the workshop, the average score increased by 4.8 to an overall average of 8.9, indicating most participants enjoyed the workshop (Figure 1A, Post-Test). Similarly, the average initial score for believing the workshop would help improve communication skills was 5.2 (Figure 1B, Pre-Test). Post-workshop, the average score increased by 4.8 points to an average of 9.9 (Figure 1B, Post-Test). The belief that the workshop would increase networking skills increased by 5.6 points; the average pre-test score was 3.9, while the average post-test score was 9.5 (Figure 1C). More so, initially, participants did not strongly believe the workshop would underscore the importance of having a support team, giving an average score of 3.6 (Figure 1D, Pre-Test). Following the workshop, this score rose by 5.7 to an average of 9.3 (Figure 1D, Post-Test). The metrics measured showed that, on average, mentees found the workshop informative and beneficial to developing their networking, communication, and collaboration skills (Figure 1A-D, Post-Test).

These workshops allow for trainees to explore concepts about adequate mentorship (Figure 1A-D). It is possible to interpret the data that initially students did not see the benefit of the workshop because they may have felt it was not specifically targeting them or irrelevant to their goals of developing a career at the undergraduate level (Figure 1A-D, Pre-Test). It is important to highlight that the students did achieve a sense of award from gleaning new information about mentorship. Taken together, these results suggest that career development workshops focused on mentorship may have a large impact on student development and performance level at the undergraduate level (Figure 1A-D).

## Discussion

Mentoring is an important aspect of a mentee’s career, especially those from URM groups (Hinton Jr *et al*. 2020a, 2020b). Taken together, the data from the questionnaire highlights the need for more career development opportunities focused on mentorship. This workshop also provided an opportunity for self-reflection for students to understand the importance of mentors and how they may be an important asset to achieving career goals. This workshop also provided a unique understanding of different mentoring practices, and which may be most effective or ineffective in a mentee-mentor relationship. Initially, enthusiasm for this type of program was low and students thought career development workshops were not essential to their development (Figure 1A-D, Pre-Test). However, the post-test results suggest that the students found this workshop offered a robust set of strategies and tools to use in their career development and an understanding of what type of mentor-mentee relationships they may need (Figure 1A-D, Post-Test).

Although our sample size was small, the data suggest that career development workshops are important for career advancement. We would further speculate that career advancement can be done within mentee-mentor relationships, as well as, through skill and knowledge-building workshops. We also suggest that students that experience this workshop can improve their overall skill set and help build an understanding of the need for introspection and evaluation of what may be helpful in their career advancement.

However, our dataset does not reflect a large stratification of participants by race and ethnicity, age, or sex. We suggest these workshops be given in other languages based on institutional demographics to effectively communicate the importance of career development to non-native speakers. Additionally, our study participants, although involved in STEM fields, may not represent the entire student-body population. Thus, we suggest that this workshop and others be used to create a series to further enrich undergraduate career development across a wide variety of demographics. Equally, we suggest that workshops like these continue to be a resource to individuals that do not have access to career development opportunities. These workshops should be open access for others to disseminate the information and help broaden the true participation and motivation needed to pursue a STEM career. Furthermore, additional study is needed to identify additional areas that may aid in student success.

## Availability of data and materials

A PowerPoint presentation of the workshop is available in English and Spanish upon request. Survey data may be made available upon reasonable request.

## Acknowledgements

We thank the 24 students who participated in our survey.

## Ethics Declaration, Project Title

Promoting Engagement in science for underrepresented Ethnic and Racial minorities (P.E.E.R), 21-MortonD-HSR-SOM-01, Kaiser Foundation Research Institute FWA: FWA00002344

## Ethics Approval and consent to participate

Yes

## Consent for publication

Yes

